# Exact Sketch-Based Read Mapping

**DOI:** 10.1101/2023.06.21.545862

**Authors:** Tizian Schulz, Paul Medvedev

## Abstract

Given a sequencing read, the broad goal of read mapping is to find the location(s) in the reference genome that have a “similar sequence”. Traditionally, “similar sequence” was defined as having a high alignment score and read mappers were viewed as heuristic solutions to this well-defined problem. For sketch-based mappers, however, there has not been a problem formulation to capture what problem an exact sketch-based mapping algorithm should solve. Moreover, there is no sketch-based method that can find all possible mapping positions for a read above a certain score threshold.

In this paper, we formulate the problem of read mapping at the level of sequence sketches. We give an exact dynamic programming algorithm that finds all hits above a given similarity threshold. It runs in 𝒪 (|*t*| + |*p*| + *ℓ*^2^) time and Θ(*ℓ*^2^) space, where |*t*| is the number of *k*-mers inside the sketch of the reference, |*p*| is the number of *k*-mers inside the read’s sketch and *ℓ* is the number of times that *k*-mers from the pattern sketch occur in the sketch of the text. We evaluate our algorithm’s performance in mapping long reads to the T2T assembly of human chromosome Y, where ampliconic regions make it desirable to find all good mapping positions. For an equivalent level of precision as minimap2, the recall of our algorithm is 0.88, compared to only 0.76 of minimap2.

**2012 ACM Subject Classification:** Applied computing → Computational biology

## 1 Introduction

Read mapping continues to be one of the most fundamental problems in bioinformatics. Given a read, the broad goal is to find the location(s) in the reference genome that have a “similar sequence”. Traditionally, “similar sequence” was defined as having a high alignment score and read mappers were viewed as heuristic solutions to this well-defined problem. However, the last few years has seen the community embrace sketch-based mapping methods, best exemplified by minimap2 [11] (see [16] for a survey). These read mappers work not on the original sequences themselves but on their sketches, e.g. the minimizer sketch. As a result, it is no longer clear which exact problem they are trying to solve, as the definition using an alignment score is no longer directly relevant. To the best of our knowledge, there has not been a problem formulation to capture what problem an exact sketch-based mapping algorithm should solve.

In this work, we provide a problem formulation (Section 3) and an exact algorithm to find all hits above a given score (Section 6). More formally, we consider the problem of taking a sketch *t* of a text *T* and a sketch *p* of a query *P* and identifying all sub-sequences of *t* that match *p* with a score above some threshold. A score function could for example be the weighted Jaccard index, though we explore several others in this paper (Section 4). We provide both a simulation-based and an analytical-based method for setting the score threshold (Section 5). Our algorithm runs in time 𝒪 (|*t*| + |*p*| + *ℓ*^2^) and space Θ(*ℓ*^2^), where *ℓ* is the number of times that *k*-mers from *p* occur in *t*.

Other sketch-based mappers are heuristic: they typically find matching elements between the reference and the read sketches (i.e. anchors) and extend these into maps using chaining [16]. Our algorithm is more resource intensive than these heuristics, as is typical for exact algorithms. However, a problem formulation and an exact algorithm gives several long-term benefits. First, the exact algorithm could be used in place of a greedy heuristic when the input size is not too large. Second, the formulation can spur development of exact algorithms that are optimized for speed and could thus become competitive with heuristics. Third, the formulation could be used to find the most effective score functions, which can then guide the design of better heuristics. Finally, our exact algorithm can return all hits with a score above a threshold, rather than just the best mapping(s). This is important for tasks such as the detection of copy number variation [12] or detecting variation in multi-copy gene families [1].

We evaluate our algorithm (called eskemap), using simulated long reads from the T2T human Y chromosome (Section 7). For the same level of precision, the recall of eskemap is 0.88, compared to 0.76 of minimap2. This illustrates the power of eskemap as a method to recover more of the correct hits than a heuristic method. We also compare against Winnowmap2 [10] and edlib [18], which give lower recall but higher precision than eskemap.

## 2 Preliminaries

### Sequences

Let *t* be a sequence of elements (e.g. *k*-mers) that may contain duplicates. We let |*t*| denote the length of the sequence, and we let *t*[*i*] refer to the *i*-th element in *t*, with *t*[0] being the first element. For 0 ≤ *i* ≤ *j <* |*t*|, let *t*[*i, j*] represent the subsequence (*t*[*i*], *t*[*i* + 1], …, *t*[*j*]). The set of elements in *t* is denoted by 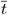, e.g. if *t* = (ACG, TTT, ACG) then *t* = {`ACG, TTT`}. We let occ(*x, t*) represent the number of occurrences of an element *x* in *t*, e.g. occ(`ACG`, *t*) = 2.

### Sketch

Let *T* be a string and let *t* be the sequence of *k*-mers appearing in *T*. Note that *t* is a sequence of DNA sequences. For example, if *T* = `ACGAC` and *k* = 2, then *t* = (AC, CG, GA, AC). For the purposes of this paper, a *sketch* of *T* is simply a subsequence of *t*, e.g. (`AC, GA`). This type of sketch could for example be a minimizer sketch [15, 17], a syncmer sketch [6], or a FracMinHash sketch [9, 7].

### Scoring Scheme

A *scoring scheme* (sc, thr) is a pair of functions: the score function and the threshold function. The *score function* sc is a function that takes as input a pair of non-empty sketches and outputs a real number, intuitively representing the degree of similarity. We assume it is symmetric, i.e. sc(*p, s*) = sc(*s, p*) for all sketches *p* and *s*. If the score function has a parameter, then we write sc(*s, p*; *θ*), where *θ* is a vector of parameter values. The *threshold function* thr takes the length of a sketch and returns a score cutoff threshold, i.e. scores below this threshold are not considered similar. Note that the scoring scheme is not allowed to depend on the underlying nucleotide sequences besides what is captured in the sketch.

### Miscellenous

We use *U*_*k*_ to denote the universe of all *k*-mers. Given two sequences *p* and *s*, the *weighted Jaccard* is defined as 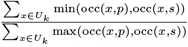. It is 0 when *s* and *p* do not share any elements, 1 when *s* is a permutation of *p*, and strictly between 0 and 1 otherwise. The weighted Jaccard is a natural extension of Jaccard similarity that accounts for multi-occurring elements.

## 3 Problem Definition

In this section, we first motivate and then define the Sketch Read Mapping Problem. Fix a scoring scheme (sc, thr). Let *p* and *t* be two sketches, which we refer to as the pattern and the text, respectively. Define a *candidate mapping* as a subinterval *t*[*a, b*] of *t*. A naive problem definition would ask to return all candidate mappings with sc(*p, t*[*a, b*]) ≥ thr(|*p*|).^1^ However, a lower-scoring candidate mapping could contain a higher-scoring candidate mapping as a subinterval, with both scores above the threshold. This may arise due to a large candidate mapping containing a more conserved small candidate mapping, in which case both candidate mappings are of interest. But it may also arise spuriously, as a candidate mapping with a score sufficiently higher than thr(|*p*|) can be extended with non-shared *k*-mers that decrease the score but not below the threshold.

To eliminate most of these spurious cases, we say that a candidate mapping *t*[*a, b*] is *reasonable* if and only if for *x* ∈ {*t*[*a*], *t*[*b*]}, occ(*x, t*[*a, b*]) ≤ occ(*x, p*). In other words, a reasonable candidate mapping must start and end with a *k*-mer that has a match in the pattern. We also naturally do not wish to report a candidate mapping that is a subinterval of a longer candidate mapping with a larger score. Formally, we call a candidate mapping *t*[*a, b*] *maximal* if there does not exist a candidate mapping *t*[*a*^*′*^, *b*^*′*^], with *a*^*′*^ ≤ *a* ≤ *b* ≤ *b*^*′*^ and sc(*t*[*a*^*′*^, *b*^*′*^], *p*) *>* sc(*t*[*a, b*], *p*). We can now formally define *t*[*a, b*] to be a *final mapping* if it is both maximal and reasonable and sc(*t*[*a, b*], *p*) ≥ thr(|*p*|). The *Sketch Read Mapping Problem* is then to report all final mappings. We now restate the problem in a succinct manner:

### ▶ Definition 1 (Sketch Read Mapping Problem).

*Given a pattern sketch p, a text sketch t, a score function sc, and a threshold function thr, the Sketch Read Mapping Problem is to find all* 0 ≤ *a* ≤ *b <* |*t*| *such that*

▪ *sc*(*p, t*[*a, b*]) ≥ *thr*(|*p*|),
▪ *occ*(*t*[*a*], *t*[*a, b*]) ≤ *occ*(*t*[*a*], *p*),
▪ *occ*(*t*[*b*], *t*[*a, b*]) ≤ *occ*(*t*[*b*], *p*),
▪ *there does not exist a*^*′*^ ≤ *a* ≤ *b* ≤ *b*^*′*^ *such that sc*(*t*[*a*^*′*^, *b*^*′*^], *p*) *> sc*(*t*[*a, b*], *p*), *i*.*e. t*[*a, b*] *is maximal*.

## 4 Score Function

In this section, we explore the design space of score functions and fix two score functions for the rest of the paper. Let *p* be the sketch of the pattern and let *s* be a continuous subsequence of the sketch of the text *t*, i.e. *s* = *t*[*a, b*] for some *a* ≤ *b*. For example if *p* = (ACT, GTA, TAC) and *t* = (AAC, ACT, CCT, GTA), we could have *s* = *t*[1, 3] = (ACT, CCT, GTA). In the context of the Sketch Read Mapping Problem, *p* is fixed and *s* varies. Therefore, while the score function is symmetric, the threshold function sets the score threshold as a function of |*p*|. Since *p* is fixed, the threshold is a single number in the context of a single problem instance. In the following, we exclusively consider score functions that calculate the similarity of *s* and *p* by ignoring the order of *k*-mers inside the sketches. Taking *k*-mer order into account would likely make it more complex to compute scores, while not necessarily giving better results on real data. However, score functions that do take order into account are possible and could provide better accuracy in some cases.

A good score function should reflect the number of *k*-mers shared between *s* and *p*. For a given *k*-mer *x*, we define

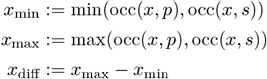

Intuitively, *x* occurs a certain number of times in *p* and a certain number of times in *s*; we let *x*_min_ be the smaller of these two numbers and *x*_max_ be the larger of these two numbers. Similarly, *x*_diff_ is the absolute difference between how often *x* occurs in *p* and *s*. We say that the number of *shared* occurrences is 2*x*_min_ and the number of *non-shared* occurrences is *x*_diff_. These quantities are governed by the relationships

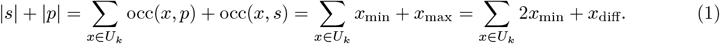

A good score function should be (1) increasing with respect to the number of shared occurences and (2) decreasing with respect to the number of non-shared occurences. There are many candidate score functions within this space. The first score function we consider is the weighted Jaccard. Formally,

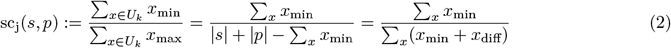

The above formula includes first the definition but then two algebraically equivalent versions of it, derived using Eq. 1. The weighted Jaccard has the two desired properties of a score function and is a well-known similarity score. However, it has two limitations. First, the use of a ratio makes it challenging to analyze probabilistically, as is the case with the non-weighted Jaccard [3]. Second, it does not offer a tuning parameter which would control the relative benefit of a shared occurence to the cost of a non-shared occurence. We therefore consider another score function, parameterized by a real-valued tuning parameter *w >* 0:

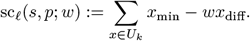

It is sometimes more useful to use an equivalent formulation, obtained using Eq. 1:

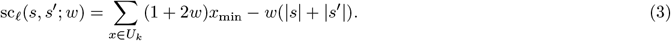

As with the weighted Jaccard, sc_*ℓ*_ has the two desired properties of a score function. But, unlike the weighted Jaccard, it is linear and contains a tuning parameter *w*.

To understand how score functions relate to each other, we introduce the notion of domination and equivalence. Informally, a score function sc_1_ dominates another score function sc_2_ when sc_1_ can always recover the separation between good and bad scores that sc_2_ can. In this case, the solution obtained using sc_2_ can always be obtained by using sc_1_ instead. Formally, let sc_1_ and sc_2_ be two score functions, parameterized by *θ*_1_ and *θ*_2_, respectively. We say that sc_1_ *dominates* sc_2_ if and only if for any parameterization *θ*_2_, threshold function thr_2_, and pattern sketch *p* there exist a *θ*_1_ and thr_1_ such that, for all sequences *s*, we have that sc_2_(*s, p*; *θ*_2_) ≥ thr_2_(|*p*|) if and only if sc_1_(*s, p*; *θ*_1_) ≥ thr_1_(|*p*|). Furthermore, sc_1_ dominates sc_2_ *strictly* if and only if the opposite does not hold, i.e. sc_2_ does not dominate sc_1_. Otherwise, sc_1_ and sc_2_ are said to be *equivalent*, i.e. if and only if each one dominates the other.

We can now precisely state the relationship between sc_*ℓ*_ and sc_j_, i.e. that sc_*ℓ*_ strictly dominates sc_j_. In other words, any solution to the Sketch Read Mapping Problem that is obtained by sc_j_ can also be obtained by sc_*ℓ*_, but not vice-versa. Formally,

### ▶ Theorem 2.

*sc*_*ℓ*_ *stricly dominates the weighted Jaccard score function sc*_*j*_.

Proof. We start by proving that sc_*ℓ*_ dominates sc_j_. Let *p* be a pattern sketch and let thr_*j*_ be the threshold function associated with sc_j_. We will use the shorthand *t* = thr_*j*_ (|*p*|). First, consider the case that *t* < 1.Let 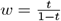 and let thr_*ℓ*_ evaluate to zero for all inputs. Let *s* be any sketch. The following is a series of equivalent transformations that proves domination.

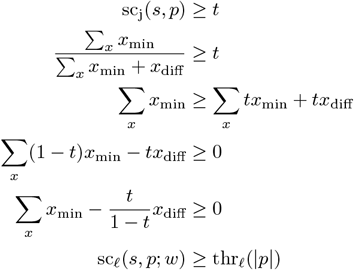

Next, consider the case *t >* 1. In this case, for all *s*, sc_j_(*s, p*) *< t*, since the weighted Jaccard can never exceed one. Observe that sc_*ℓ*_(*s, p*; *w*) ≤ |*p*| for any non-negative *w*. Therefore, we can set thr_*ℓ*_(|*p*|) = |*p*| + 1 and let *w* be any non-negative number, guaranteeing that for all *s*, sc_*ℓ*_(*s, p*; *w*) *<* thr_*ℓ*_(|*p*|).

Finally consider the case that *t* = 1. Then, sc_j_(*s, p*) ≥ *t* if and only if *s* and *p* are permutations of each other, i.e. *x*_diff_ = 0 for all *x*. Setting thr_*ℓ*_(|*p*|) = |*p*| and letting *w* be any strictly positive number guarantees that sc_*ℓ*_(*s, p*; *w*) ≥ thr_*ℓ*_(|*p*|) if and only if *s* and *p* are permutations of each other.

To prove that sc_*ℓ*_ is not dominated by sc_j_, we fix *w* = 1 (though any value could be used) and give a counter example family to show that sc_j_ cannot recover the separation that sc_*ℓ*_ can. Pick an integer *i* ≥ 1 to control the size of the counterexample. Let *p* be a pattern sketch of length 4*i* consisting of arbitrary *k*-mers. We construct two sketches, *s*_1_ and *s*_2_. The sequence *s*_1_ is an arbitrary subsequence of *p* of length *i*. Observe that 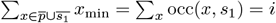. The sequence *s*_2_ is *p* appened with arbitrary *k*-mers to get a length 12*i*. Observe that 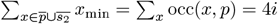. Using Eq. 3 for sc_*ℓ*_ and Eq. 2 for sc_j_,

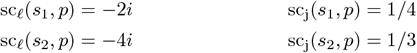

Under sc_*ℓ*_, *s*_1_ has a higher score, while under sc_j_, *s*_2_ has a higher score. If thr_*ℓ*_ is set to accept *s*_1_ but not *s*_2_ (e.g. thr_*ℓ*_ = −3*i*), then it is impossible to set thr_*j*_ to achieve the same effect. In other words, since sc_j_(*s*_2_) *>* sc_j_(*s*_1_), any threshold that accepts *s*_1_ must also accept *s*_2_.

Next, we show that many other natural score functions are equivalent to sc_*ℓ*_. Consider the following score functions:

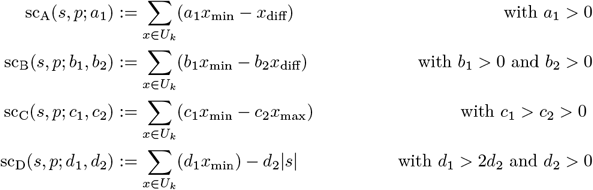

The conditions on the parameters are there to enforce the two desired properties of a score function. Each of these score functions is natural in its own way, e.g. sc_A_ is similar to sc_*ℓ*_ but places the weight on *x*_min_ rather than on *x*_diff_. One could also have two separate weights, as in the score sc_B_. One could then replace *x*_diff_ with *x*_max_, as in sc_C_, which is the straightforward reformulation of the weighted Jaccard score as a difference instead of a ratio. Or one could replace *x*_diff_ with the length of *s*, as in sc_D_. The following theorem shows that the versions turn out to be equivalent to sc_*ℓ*_ and to each other. The proof is a straightforward algebraic manipulation and is left for the appendix.

### ▶ Theorem 3.

*The score functions sc*_*ℓ*_, *sc*_*A*_, *sc*_*B*_, *sc*_*C*_, *and sc*_*D*_ *are pairwise equivalent*.

## 5 Choosing a Threshold

In this section, we propose two ways to set the score threshold. The first is analytical (Section 5.1) and the second is with simulations (Section 5.2). The analytical approach gives a closed form formula for the expected value of the score under a mutation model. However, it only applies to the FracMinHash sketch, assumes a read has little internal homology, and does not give a confidence interval. The simulation approach can apply to any sketch but does not offer any analytical insight into the behavior of the score. The choice of approach ultimately depends on the use case.

We first need to define a generative mutation model to capture both the sequencing and evolutionary divergence process:

### ▶ Definition 4 (Mutation model).

*Let S be a circular string*^2^ *with n characters. The mutation model produces a new string S*^*′*^ *by first setting S*^*′*^ = *S and then taking the following steps:*

1. *For every* 0 ≤ *i < n, draw an action a*_*i*_ ∈ {*sub, del, unchanged*} *with probability of p*_*sub*_ *for sub, p*_*del*_ *for del, and* 1 − *p*_*sub*_ − *p*_*del*_ *for unchanged. Also, draw an insertion length b*_*i*_ *from a geometric distribution with mean p*_*ins*_^3^.
2. *Let track be a function mapping from a position in S to its corresponding position in S*^*′*^. *Initially, track*(*i*) = *i, but as we delete and add characters to S*^*′*^, *we assume that track is updated to keep track of the position of S*[*i*] *in S*^*′*^.
3. *For every i such that a*_*i*_ = *sub, replace S*^*′*^[*i*] *with one of the three nucleotides not equal to S*[*i*], *chosen uniformly at random*.
4. *For every* 0 ≤ *i < n, insert b*_*i*_ *nucleotides (chosen uniformaly at random) before S*^*′*^[*track*(*i*)].
5. *For every i such that a*_*i*_ = *del, remove S*^*′*^[*track*(*i*)] *from S*^*′*^.

### 5.1 Analytical Analysis

To derive an expected score under the mutation model, we need to specify a sketch. We will use the FracMinHash sketch [9], due its simpliticy of analysis [7].

#### ▶ Definition 5 (FracMinHash).

*Let h be a hash function that maps a k-mer to a real number between 0 and 1, inclusive. Let* 0 *< q* ≤ 1 *be a real-valued number called the sampling rate. Let S be a string. Then the* FracMinHash *sketch of S, denoted by s, is the sequence of all k-mers x of S, ordered as they appear in S, such that h*(*x*) ≤ *q*.

Consider an example with *k* = 2, *S* = CGGACGGT, and the only *k*-mers hashing to a value ≤ *q* being `CG` and `GG`. Then, *s* = (CG, GG, CG, GG).

We make an assumption, which we refer to as the *mutation-distinctness assumption*, that the mutations on *S* never create an *k*-mer that is originally in *S*. Based on previous work [4], we find this necessary to make the analysis mathematically tractable (for us). The results under this assumption become increasingly inaccurate as the read sequence contains increasingly more internal similarity. For example, reads coming from centromeres might violate this assumption. In such cases, it may be better to choose a threshold using the technique in Section 5.2.

We can now derive the expected value of the score under the mutation model and FracMinHash.

#### ▶ Theorem 6.

*Let S be a circular string and let S*^*′*^ *be generated from S under the mutation model with the mutation-distinctness assumption and with parameters p*_*sub*_, *p*_*del*_, *and p*_*ins*_. *Let s and s*^*′*^ *be the FracMinHash sketches of S and S*^*′*^, *respectively, with sampling rate q. Then, for all real-valued tuning parameters w >* 0,

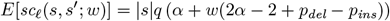

*where* 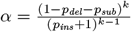.

Proof. Observe that under mutation-distinctness assumption, the number of occurrences of a *k*-mer that is in *s* can only decrease after mutation, and a *k*-mer that is newly created after mutation has an *x*_min_ of 0. Therefore, applying Eq. 3,

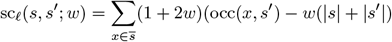

(Recall that 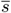 is the set of all *k*-mers in *s*.) We will first compute the score conditioned on the hash function of the sketch being fixed. Note that when *h* is fixed, then the sketch *s* becomes fixed and *s*^*′*^ becomes only a function of *S*^*′*^. By linearity of expectation,

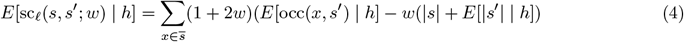

It remains to compute *E*[|*s*^*′*^| | *h*] and *E*[occ(*x, s*^*′*^) | *h*]. Observe that the number of elements in *s*^*′*^ is the number of elements in *s* minus the number of deletions plus the sum of all the insertion lengths. By linearity of expectation,

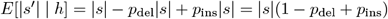

Next, consider a *k*-mer 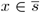 and *E*[occ(*x, s*^*′*^)]. Recall by our mutation model that no new occurrenes of *x* are introduced during the mutation process. So occ(*x, s*^*′*^) is equal to the number of occurrences of *x* in *S* that remain unaffected by mutations. Consider an occurrence of *x* in *s*. The probability that it remains is the probability that all actions on the *k* nucleotides of *x* were “unchanged” and the length of all insertions in-between the nucleotides was 0. Therefore,

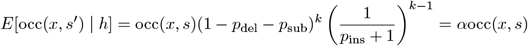

Putting it all together,

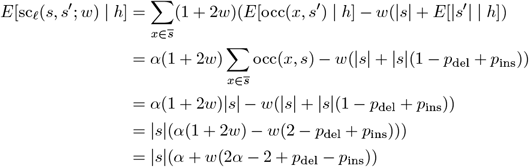

To add the sketching step, we know from [7] that the expected size of a sketch is the size of the original text times *q*. Then,

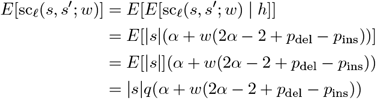

### 5.2 Simulation-Based Analysis

First, we choose the parameters of the mutation model according to the target sequence divergence between the reads and the reference caused by sequencing errors, but also due to the evolutionary distance between the reference and the organism sequenced. If one is also interested in mapping reads to homologous regions within the reference that are related more distantly, e.g. if there exist multiple copies of a gene, the mutation parameters can be increased further.

To generate a threshold for a given read length, we generate sequence pairs (*S, S*^*′*^), where *S* is a uniformly random DNA sequence of the given length and *S*^*′*^ is mutated from *S* under the above model. We then calculate the sketch of *S* and *S*^*′*^, which we call *s* and *s*^*′*^, respectively. The sketch can for example be a minimizer sketch, a syncmer sketch, or a FracMinHash sketch. We can then use the desired score function to calculate a score for each pair (*s, s*^*′*^). For a sufficiently large number of pairs, their scores will form an estimate of the underlying score distribution for sequences that evolved according to the used model. It is then possible to choose a threshold such that the desired percentage of mappings would be reported by our algorithm. For example, one could choose a threshold to cover a one sided 95% confidence interval of the score.

In order to be able to adjust thresholds according to the variable length of reads produced from a sequencing run, the whole process may be repeated several times for different lengths of *S*. Thresholds can then be interpolated dynamically for dataset reads whose lengths were not part of the simulation.

## 6 Algorithm for the Sketch Read Mapping Problem

In this section, we describe a dynamic programming algorithm for the Sketch Read Mapping Problem under both the weighted Jaccard and the linear scores (sc_j_ and sc_*ℓ*_, respectively). Let *t* be the sketch of the text, let *p* be the sketch of the pattern, let *L* be the sequence of positions in *t* that have a *k*-mer that is in 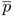, in increasing order, and let *ℓ* = |*L*|. Our algorithm takes advantage of the fact that *p* is typically much shorter than *t* and hence the number of elements of *t* that are shared with *p* is much smaller than |*t*| (i.e. *ℓ* ≪ |*t*|). In particular, it suffices to consider only candidate mappings that begin and end in positions listed in *L*, since by definition, if *t*[*a, b*] is a reasonable candidate mapping, then 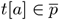 and 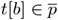.

We present our algorithm as two parts. In the first part (Section 6.1), we compute a matrix *S* with *ℓ* rows and *ℓ* columns so that *S*(*i, j*) = Σ_*x*_ min(occ(*x, p*), occ(*x, t*[*L*[*i*], *L*[*j*]]). *S* is only defined for *j* ≥ *i*. We also mark each cell of *S* as being reasonable or not. In the second part (Section 6.2), we scan through *S* and output the candidate mapping *t*[*i, j*] if and only if it is maximal and has a score above the threshold.

The reason that *S*(*i, j*) is not defined to store the score of the candidate mapping *t*[*L*[*i*], *L*[*j*]] is that the score can be computed from *S*(*i, j*) in constant time, for both sc_j_ and sc_*ℓ*_. To see this, let *x*_min_ := min(occ(*x, p*), occ(*x, t*[*L*[*i*], *L*[*j*]]). Recall that Equation (2) allows us to express sc_j_(*t*[*i, j*], *p*) as a function of Σ*x*_min_, |*p*|, and the length of the candidate mapping, i.e. *j* − *i* + 1. Similarly, we can apply Equation (1) to express sc_*ℓ*_ as

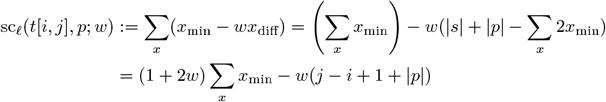

Thus, once Σ_*x*_ *x*_min_ is computed, either of the scores can be computed trivially.

### 6.1 Computing *S*

We compute *S* using dynamic programming. For the base case of the diagonal, i.e. for 0 ≤ *i < ℓ*, we can set *S*(*i, i*) = 1. Here, since we know that 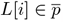, we get that the *k*-mer *t*[*L*[*i*]] occurs at least once in *p* and exactly once in *t*[*L*[*i*], *L*[*i*]]. For the general case, i.e. for 0 ≤ *i < j < ℓ*, we can define *S* using a recursive formula:

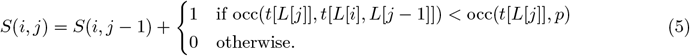

To see the correctness of this formula, observe that all the elements of *t*[*L*[*j* −1]+1, *L*[*j*]−1] are, by definition, not in 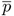 and hence their minimum occurrence value is 0. If the element *x* = *t*[*L*[*j*]] helps increase min(occ(*x, t*[*L*[*i*], *L*[*j*−1]]), occ(*x, p*)), then we increase the minimum count by one, otherwise the minimum occurrence does not increase. Furthermore, we can mark *S*(*i, j*) as being right-reasonable anytime that the top case is used and as not being right-reasonable otherwise.

To design an efficient algorithm based on Equation (5), we need two auxiliary data structures. The first is a hash table *H*_*cnt*_ that stores, for every *k*-mer in 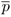, how often it occurs in *p*. A second hash table *H*_*loc*_ stores, for every *k*-mer 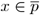, the number of locations *i* such that *t*[*L*[*i*]] = *x*.

The algorithm for computing *S* and the hash tables is given in Algorithm 1. As a first step, the *H*_cnt_ hash table is constructed via a scan through *p*. Then, the *S* matrix is filled in column-by-column using Equation (5). However, doing the check to determine which case of Equation (5) to use (i.e. to compute occ(*t*[*L*[*j*]], *t*[*L*[*i*], *L*[*j* − 1]])) would take non-constant time using a naive approach. In order to compute this in constant time, let *c*_1_ = occ(*x, t*[0, *L*[*i* − 1]]) and let *c*_2_ = occ(*x, t*[0, *L*[*j* − 1]]) and observe that occ(*x, t*[*L*[*i*], *L*[*j* − 1]]) = *c*_2_ − *c*_1_. We will now describe how to maintain *c*_2_ and *c*_1_ as we process a column of *S*, with only constant time per cell.

To compute *c*_2_, we avoid building *H*_loc_ outright and instead build *H*_loc_ at the same time as we are processing *S*, column-by-column. When processing column *j* with *x* = *L*[*j*], we start by incrementing the count of *H*_loc_[*x*] (Line 7). Let 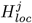 refer to *H*_loc_ right after making this insertion. Observe that 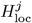 is *H*_loc_ but only containing the counts of locations up to *L*[*j*], and 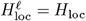. Computing *c*_2_ is trivial from 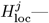 it is simply 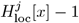 (Line 9).

To compute *c*_1_, we use the fact that when computing a column of *S*, we are processing all the rows starting from 0 up to *ℓ* − 1. We initially set *c*_1_ = 0 (Line 8) and then, for each new row *i*, we increment *c*_1_ if *t*[*L*[*i*]] = *x* (Line 17).

After *S* has been filled, we can identify which of the candidate mappings are reasonable. Observe that a candidate mapping *t*[*L*[*i*], *L*[*j*]] is reasonable if and only if *S*(*i, j*) *> S*(*i* + 1, *j*) and *S*(*i, j*) *> S*(*i, j* − 1). This can be verified by a simple pass through the matrix (Lines 22-31).

### 6.2 Computing Maximality

In the second step, we identify which of the candidate mappings in *S* are maximal. Our algorithm is shown in Algorithm 2. We traverse *S* column-by-column starting with the last column and then row-by-row, starting from the first row. While traversing *S*, we maintain a list *M* of all maximal reasonable candidate mappings above the threshold found so far and their scores. *M* has the invariant that the candidate mappings are increasingly ordered by their start positions.

To maintain the invariant that *M* is sorted by start position, we maintain a pointer *cur* to a location in *M* (Lines 6-9). At the start of a new column traversal, when the row *i* = 0, *cur* points to the start of *M*. As the row is increased, we move *cur* forward until it hits the first value in *M* with a start larger than *i*. When a new final mapping is added to *M*, we do so at *cur*, which guarantees the order invariant of *M* (Lines 14-18).

Due to the order cells in *S* are processed during our traversal, a candidate mapping *t*[*L*[*i*], *L*[*j*]] is maximal if and only if its score is larger than the score of all other final mappings in *M* with position *i*^*′*^ ≤ *i*. For a given column, since we are processing the candidate mappings in increasing order of *i*, we can simultenously maintain a running variable *maxSoF ar* that holds the maximum value in *M* up *cur* (Line 7). We can then determine if a candidate mapping is maximal by simply checking its score against *maxSoF ar*

(Line 12).

#### Algorithm 1

Part 1 of eskemap Algorithm

*Input*: two sketches *t* and *p*

*Output*: the matrix *S* and annotation of each upper diagonal cell as being reasonable or not.

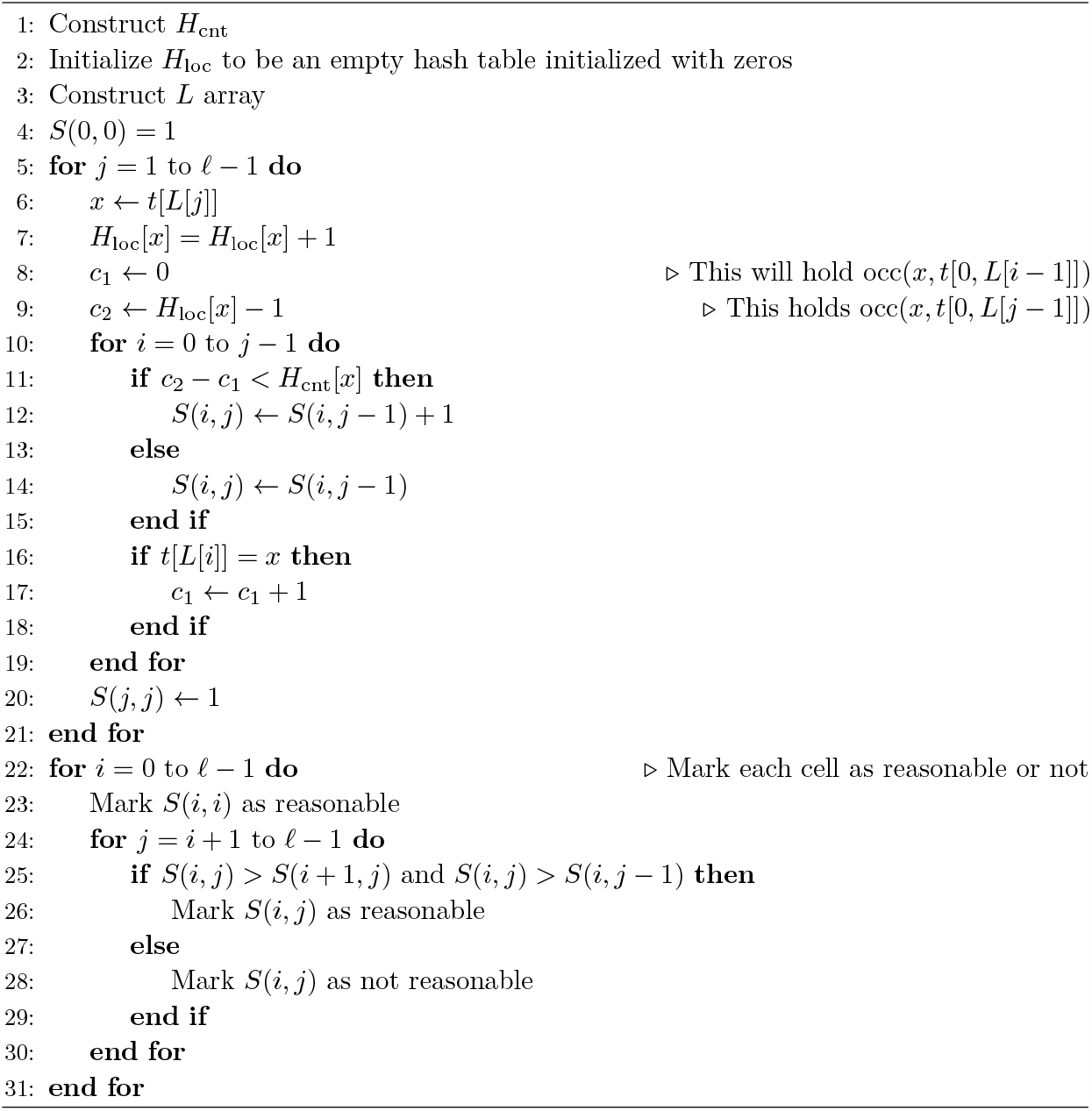

#### Algorithm 2

Part 2 of eskemap algorithm

*Input*: two sketches *t* and *p*, the matrix *S* computed by Algorithm 1, a score function, and a threshold function thr.

*Output*: all final mappings that are reasonable, maximal, and have a score of at least thr(|*p*|).

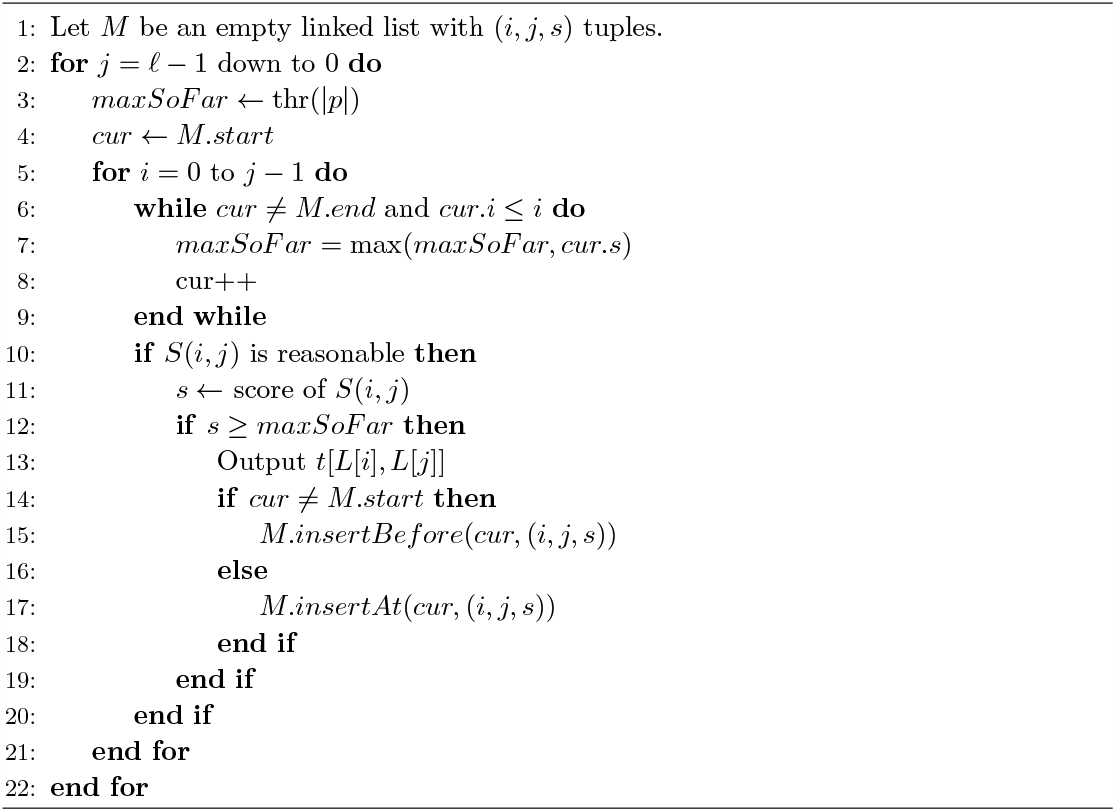

### 6.3 Runtime and Memory Analysis

The runtime for Algorithm 1 is Θ(*ℓ*^2^) + |*t*| + |*p*|. Note that the *H*_cnt_ table can be constructed in a straightforward manner in 𝒪 (|*p*|) time, assuming a hash table with constant insertion and lookup time; the *L* array is constructed in 𝒪 (|*t*|). Algorithm 2 runs two for loops with constant time internal operations, with the exception of the while loop to fast forward the *cur* pointer. The total time that for the loop is amortized to *O*(*ℓ*) for each column. Therefore, the total time for Algorithm 2 is Θ(*ℓ*^2^). This gives the total running time for our algorithm as 𝒪 (|*t*| + |*p*| + *ℓ*^2^).

The total space used by the algorithm is the sum of the space used by *S* (i.e. Θ(*ℓ*^2^)) and the space used by *H*_cn_, 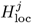, and *L*. The *H*_cnt_ table stores |*p*| integers with values up to |*p*|. However, notice that when |*p*| *> ℓ*, we can limit the table to only store *k*-mers that are in 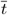, i.e. only *ℓ k*-mers. We can also replace integer values greater than *ℓ* with *ℓ*, as it would not affect the algorithm. Therefore, the *H*_cnt_ table uses 𝒪 (*ℓ* log *ℓ*) space. The 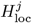 table stores at most *ℓ* entries with values at most *ℓ* and therefore takes Θ(*ℓ* log *ℓ*) space. Thus our algorithm uses a total of Θ(*ℓ*^2^) space.

## 7 Results

We implemented the eskemap algorithm described in Section 6 using sc_*ℓ*_ as score function and compared it to other methods in a read mapping scenario. For better comparability, we implemented it with the exact same minimizer sketching approach as used by minimap2. Source code of our implementation as well as a detailed documentation of our comparison described below including exact program calls is available from https://github.com/medvedevgroup/eskemap.

### 7.1 Datasets

For our evaluation, we used the T2T reference assembly of human chromosome Y (T2T-CHM13v2.0) [13]. The chromosome contains many ampliconic regions with duplicated genes from several gene families. Identifying a single best hit for reads from such regions is not helpful and instead it is necessary to find all good mappings [5]. Such a reference poses a challenge to heuristic algorithms and presents an opportunity for an all-hits mapper like eskemap to be worth the added compute.

We simulated a read dataset on this assembly imitating characteristics of a PacBio Hifi sequencing run [8]. For each read, we randomly determined its length *r* according to a gamma distribution with a 9000bp mean and a standard deviation of 7000bp. Afterwards, a random integer *i* ∈ [1, *n* − *r* + 1] was drawn as the read’s start position, where *n* refers to the length of the chromosome. Sequencing errors were simulated by introducing mutations into each read’s sequence using the mutation model described in Definition 4 and a total mutation rate of 0.2% distributed with a ratio of 6:50:54 between substitution/insertion/deletion, as suggested in [14]. Aiming for a sequencing depth of 10x, we simulated 69401 reads.

The T2T assembly of the human chromosome Y contains long centromeric and telomeric regions which consist of short tandem and higher order repeats. Mapping reads in such regions results in thousands of hits that are meaningless for many downstream analyses and significantly increases the runtime of mapping. Therefore, we excluded all reads from the initially simulated set which could be aligned to more than 20 different, non-overlapping positions using edlib (see below). After filtering, a set of 32295 reads remained.

### 7.2 Tools

We compared eskemap to two other sketch-based approaches and an exact alignment approach. The sketch-based approaches were minimap2 (version 2.24-r1122) and Winnowmap2 (version 2.03), run using default parameters. In order to be able to compare our results also to an exact, alignment-based mapping approach, we used the C/C++ library of Edlib [18] (version 1.2.7) to implement a small script that finds all non-overlapping substrings of the reference sequence a read could be aligned to with an edit distance of at most *T*. We tried values *T* ∈ {0.01*r*, 0.02*r*, 0.03*r*}, where recall that *r* is the read length. We refer to this script as simply *edlib*.

For eskemap, we aimed to make the results as comparable as possible to minimap2. We therefore used a minimizer sketch with the same *k*-mer and window size as minimap2 (*k* = 15, *w* = 10). However, we excluded minimizers that occurred *>* 100 times inside the reference sketch, to limit the *O*(*ℓ*^2^) memory use of eskemap, even as this exclusion may potentially hurt eskemap’s accuracy. We used the default *w* = 1 as the tuning parameter in the linear score. To set the score threshold, we used the dynamic procedure described in Section 5.2. In particular, we used five different sequence lengths for simulations and used a divergence of 1%. We used the same sequencing error profile as for read simulation. Four thresholds were then chosen so at to cover the one-sided confidence interval of 70%, 80%, 90%, and 95%, respectively.

### 7.3 Accuracy Measure

We compared the reference substrings corresponding to each reported mapping location of any tool to the mapped read’s sequence using BLAST [2]. If a pairwise comparison of both sequences resulted either in a single BLAST hit with an E-value not exceeding 0.01^4^ and covering at least 90% of the substring or the read sequence or if a set of non-overlapping BLAST hits was found of which none had an E-value above 0.01 and their lengths summed up to at least 90% of either the reference substring’s or the read sequence’s length, we considered the reference mapping location as homologous.

For each read, we combine all the homologous reference substrings found across all tools into a ground truth set for that read. We then measure the accuracy of a mapping as follows. We determined for each *k*-mer of the reference sequence’s sketch whether it is either a *true positive* (TP), *false positive* (FP), *true negative* (TN) or *false negative* (FN). A *k*-mer was considered a TP if it was covered by both a mapping and a ground truth substring. It was considered a FP if it was covered by a mapping, but not by any ground truth substring. Conversely, it was considered a TN if it was covered by neither a mapping nor a ground truth substring and considered a FN if it was covered by a substring of the ground truth exclusively. The determination was performed for each read independently and results were accumulated per tool to calculate precision and recall measures.

### 7.4 Accuracy Results

The precision and recall of the various tools is shown in Figure 2. The most controlled comparison can be made with respect to minimap2, since the sketch used by eskemap is a subset of the one used by minimap2. At a score threshold corresponding to 70% recovery, eskemap achieves the same precision (0.999) as minimap2. However, the recall of eskemap is 0.88, compared to 0.76 of minimap2. This illustrates the potential of eskemap as a method to recover more of the correct hits than a heuristic method. More generally, eskemap achieves a recall around 90%, while all other tools have a recall of at most 76%. However, both edlib and Winnowmap2 achieve a slightly higher precision (by 0.001).

**Figure 1.**
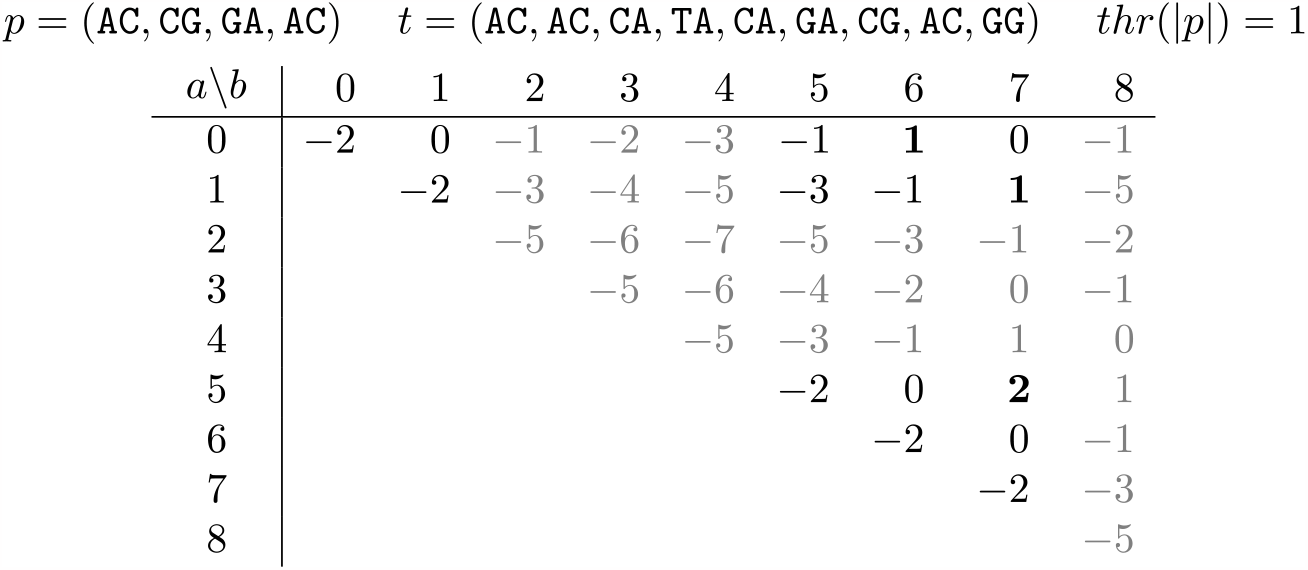
An example of the Sketch Read Mapping Problem. We show all candidate mappings *t*[*a, b*] for a given pattern *p* and a text *t*. Each candidate mapping is represented by its score calculated using *sc*_*ℓ*_(*p, t*[*a, b*]; 1) (see Section 4). Reasonable candidate mappings are shown in black (rather than gray) and final mappings are further bolded.

**Figure 2.**
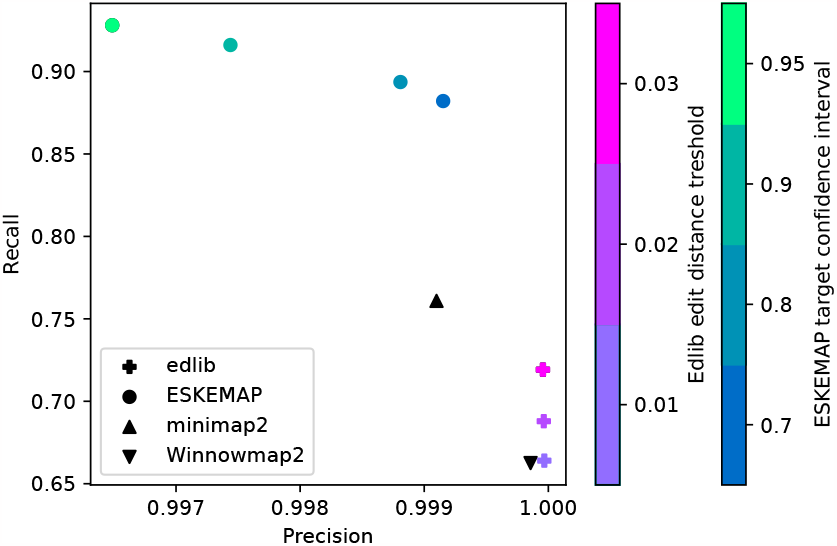
Mapping accuracies of all tools. For edlib, the color of the cross encodes the various edit distance thresholds (0.01, 0.02, 0.03). For eskemap, the color of the circles indicate the score threshold used, in terms of the target confidence interval used (0.7, 0.8, 0.9, 0.95). The ground truth is determined by combining the mappings from all tools and filtering out those with bad BLAST scores. The most lenient thresholds for edlib and eskemap were used.

### 7.5 Time and Memory Results

We compared the runtimes and memory usage of all sketch-based methods (Table 1). Calculations were performed on a virtual machine with 28 cores and 256 GB of RAM. We did not include edlib in this alignment since, as an exact alignment-based method, it took much longer to complete (i.e. running highly parallelized on many days on a system with many cores). We see that both heuristics are significantly faster than our exact algorithm. However, they also find many fewer mapping positions per read. E.g., only one mapping position is reported for 67% and 75% of all reads by minimap2 and Winnowmap2, respectively. In comparison, eskemap finds more than one mapping position for almost every second read (49%). When the runtime is normalized per output mapping, eskemap is actually more than an order of magnitude faster than the other tools.

**Table 1.**
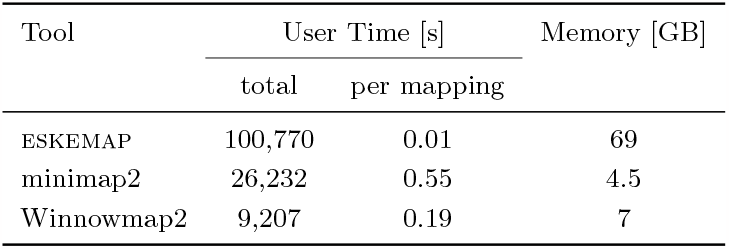
Runtime and memory usage comparison of all sketch-based methods. The tools were called to map 32295 simulated PacBio Hifi sequencing reads on the T2T assembly of human chromosome Y. Runtimes are shown both as total values and normalized by the number of reported mapping positions.

The memory usage of eskemap is dominated by the size of *S*. In particular, the highest value of *ℓ* was 185, 702 and a matrix with dimensions *ℓ × ℓ* that stores a 4-byte value in the upper diagonal takes 69GB, which corresponds to the peak reported in Table 1. As expected, the memory use depends on the repetitiveness of the text and on the sketching scheme used.

## 8 Conclusion

In this work, we formally defined the Sketch Read Mapping Problem, i.e. to find all positions inside a reference sketch with a certain minimum similarity to a read sketch under a given similarity score function. We also proposed an exact dynamic programming algorithm called ESKEMAP to solve the problem, running in 𝒪 (|*t*| + |*p*| + *ℓ*^2^) time and Θ(*ℓ*^2^) space. We evaluated eskemap’s performance by mapping a simulated long read dataset to the T2T assembly of human chromosome Y and found it to have a superior recall for a similar level of precision compared to minimap2, while offering precision/recall tradeoffs compared with edlib or Winnowmap2.

A clear drawback of eskemap remains its high memory demand for storing the dynamic programming matrix. If many *k*-mers from a read’s sketch occur frequently inside the sketch of the reference sequence, its quadratic dependence on the number of shared *k*-mers becomes a bottleneck. It may be possible to modify the algorithm to store only the recently calculated column, but that would require a novel way to perform the maximality check of Algorithm 2. In order to further improve on eskemap’s runtime, a strategy could be to develop filters that prune the result’s search space. This could be established, e.g., by terminating score calculations for a column once its clear an optimal solution would not make use of the rest of that column. Our prototype implementation of eskemap would also benefit from additional engineering of the code base, potentially leading to substantial improvements of runtime and memory in practice.

Having an exact sketch-based mapping algorithm at hand also opens the door for the exploration of novel score functions to determine sequence similarity on the level of sketches. Using our algorithm, combinations of different sketching approaches and score functions may be easily tested. Eventually, this may lead to a better understanding of which sketching methods and similarity measures are most efficient considering sequences with certain properties like high repetitiveness or evolutionary distance.

## Supplementary Material

*Software (Source Code)*: https://github.com/medvedevgroup/eskemap

## Acknowledgements

This work was supported by the BMBF-funded de.NBI Cloud within the German Network for Bioinformatics Infrastructure (de.NBI) (031A532B, 031A533A, 031A533B, 031A534A, 031A535A, 031A537A, 031A537B, 031A537C, 031A537D, 031A538A).

## 9 Proofs

### Proof of Theorem 3.

Observe that domination is a transitive property, i.e. if sc_1_ dominates sc_2_ and sc_2_ dominates sc_3_, then sc_1_ dominates sc_3_. To prove equivalence, we will prove the following circular chain of domination: sc_*ℓ*_ ← sc_B_ ← sc_C_ ← sc_D_ ← sc_A_ ← sc_*ℓ*_.

First, observe that sc_B_ trivially dominates sc_*ℓ*_ by keeping the threshold function the same and setting *b*_1_ = 1 and *b*_2_ = *w*.

Next, we prove that sc_C_ dominates sc_B_. Let *p* be a pattern and let *t* = thr_B_(|*p*|). Set thr_C_ = thr_B_ and *c*_1_ = *b*_1_ + *b*_2_ and *c*_2_ = *b*_2_. Then, for all *s*, the following series of equivalent transformations proves that sc_C_ dominates sc_B_.

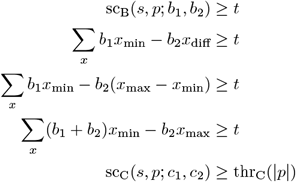

Next, we prove that sc_D_ dominates sc_C_. Let *p* be a pattern and let *t* = thr_C_(|*p*|). Set *d*_1_ = *c*_1_ + *c*_2_, *d*_2_ = *c*_2_, and thr_D_(*i*) = thr_C_(*i*) + *ic*_2_. Then, for all *s*, the following series of equivalent transformations proves that sc_D_ dominates sc_C_.

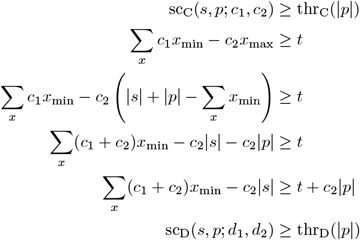

Next, we prove that sc_A_ dominates sc_D_. Let *p* be a pattern and let *t* = thr_D_(|*p*|). Set 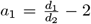 and 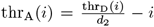. Then, for all *s*, the following series of equivalent transformations proves that sc_D_ dominates sc_C_.

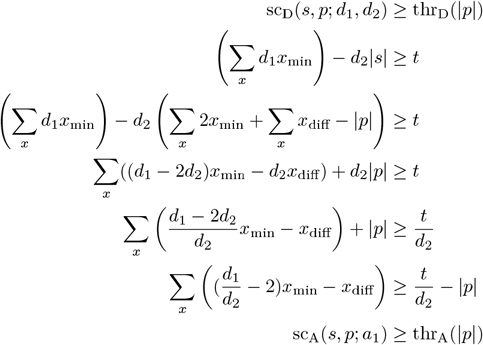

Finally, we prove that sc_*ℓ*_ dominates sc_A_. Let *p* be a pattern and let *t* = thr_A_(|*p*|). Set 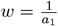 and 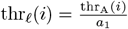. Then, for all *s*, the following series of equivalent transformations proves that sc_*ℓ*_ dominates sc_A_.

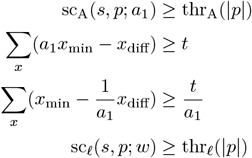

## Footnotes

1 Notice that in this framing, the threshold is not a single parameter but can vary depending on the read length. This gives flexibility to the scoring function, since the scores of candidate mappings of reads of different lengths do not need to be comparable to each other. Moreover, computing the threshold value is not a challenge since it needs to be computed just once for each read.

2 We assume the string is circular to avoid edge cases in the analysis but, for long enough strings, this assumption is unlikely to effect the accuracy of the results.

3 Here, a geometric distribution is the number of failures before the first success of a Bernoulli trial. This geometric distribution has parameter 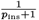.

4 In order to ensure robustness of results, BLAST runs were also repeated for E-value thresholds of 0.005 and 0.001 causing only neglectable differences for subsequent analyses.

